# Inexpensive monitoring of flying insect activity and abundance using wildlife cameras

**DOI:** 10.1101/2021.08.24.457487

**Authors:** Jesse R A Wallace, Therese Reber, Brendan Beaton, David Dreyer, Eric J Warrant

**Affiliations:** Research School of Biology, Australian National University; Lund Vision Group, Department of Biology, Lund University, Sweden

**Keywords:** Bogong moth, camfi, insect flight behaviour, insect monitoring, object detection, trail camera, wildlife camera, wingbeat frequency

## Abstract

1. The ability to measure flying insect activity and abundance is important for ecologists, conservationists and agronomists alike. However, existing methods are laborious and produce data with low temporal resolution (e.g. trapping and direct observation), or are expensive, technically complex, and require vehicle access to field sites (e.g. radar and lidar entomology).
2. We propose a method called “camfi” for long-term non-invasive monitoring of the activity and abundance of low-flying insects using images obtained from inexpensive wildlife cameras, which retail for under USD$100 and are simple to operate. We show that in certain circumstances, this method facilitates measurement of wingbeat frequency, a diagnostic parameter for species identification. To increase usefulness of our method for very large monitoring programs, we have developed and implemented a tool for automatic detection and annotation of flying insect targets based on the popular Mask R-CNN framework. This tool can be trained to detect and annotate insects in a few hours, taking advantage of transfer learning.
3. We demonstrate the utility of the method by measuring activity levels and wingbeat frequencies in Australian Bogong moths *Agrotis infusa* in the Snowy Mountains of New South Wales, and find that these moths have log-normally distributed wingbeat frequencies (mean = 49.4 Hz, std = 5.25 Hz), undertake dusk flights in large numbers, and that the intensity of their dusk flights is modulated by daily weather factors. Validation of our tool for automatic image annotation gives baseline performance metrics for comparisons with future annotation models. The tool performs well on our test set, and produces annotations which can be easily modified by hand if required. Training completed in less than 2 h on a single machine, and inference took on average 1.15 s per image on a laptop.
4. Our method will prove invaluable for ongoing efforts to understand the behaviour and ecology of the iconic Bogong moth, and can easily be adapted to other flying insects. The method is particularly suited to studies on low-flying insects in remote areas, and is suitable for very large-scale monitoring programs, or programs with relatively low budgets.

## 1 Introduction

The ability to measure flying insect activity and abundance is important for ecologists, conservationists and agronomists alike. Traditionally, this is done using tedious and invasive methods including nets (e.g. Drake & Farrow, 1985), window traps (e.g. Knuff et al., 2019), light traps (e.g. Beck et al., 2006; Infusino et al., 2017), and pheromone traps (e.g. Athanassiou et al., 2004; Laurent & Frérot, 2007), with the latter being favoured by agronomists for its specificity. The WWII development of radar led to the introduction of radar ornithology (Eastwood, 1967; Gauthreaux Jr & Belser, 2003), and ultimately radar entomology (Drake & Reynolds, 2012; Riley, 1989), which facilitated non-invasive remote sensing of insects flying up to a couple of kilometres above the ground, and became extremely important for understanding the scale and dynamics of insect migration (Chapman et al., 2011). More recently, entomological lidar has been introduced, which benefits from a number of advantages over radar, in particular the ability to measure insects flying close to the ground, without suffering from ground clutter (Brydegaard et al., 2017; Brydegaard & Jansson, 2019). However, both entomological radar and entomological lidar systems are relatively large (requiring vehicle access to study sites), bespoke, expensive, and require expertise to operate, reducing their utility and accessibility to field biologists.

We propose a method for long-term non-invasive monitoring of the activity and abundance of low-flying insects using inexpensive wildlife cameras, which retail for under USD$100 and are simple to operate. We show that in certain circumstances, this method facilitates the measurement of wingbeat frequency, a diagnostic parameter for species identification. We demonstrate the utility of the method by measuring activity levels and wingbeat frequencies in Australian Bogong moths *Agrotis infusa* that were photographed flying across a boulder field near Cabramurra in the Snowy Mountains of New South Wales. The Bogong moth is an important source of energy and nutrients in the fragile Australian alpine ecosystem (Green, 2011), and is a model species for studying directed nocturnal insect migration and navigation (Adden et al., 2020; Dreyer et al., 2018; Warrant et al., 2016). A dramatic drop in the population of Bogong moths has been observed in recent years (Green et al., 2021; Mansergh et al., 2019), adding it to the growing list of known invertebrate species whose populations are declining (Sánchez-Bayo & Wyckhuys, 2019). The present method will prove invaluable for ongoing efforts to understand the behaviour and ecology, and monitor the population of this iconic species. Our method can easily be adapted to other flying insects, and is particularly suited to large-scale monitoring programs with limited resources.

## 2 Methods

The methods outlined below summarise the use of wildlife cameras for monitoring flying insects, and detail specific methods employed in this study. With the exception of the camera set-up (in section 2.1 and appendix A.7), and the manual image annotation (section 2.2), each of the methods described below have been automated in our freely available software. A full practical step-by-step guide for using this method along with complete documentation of the latest version of the code is provided at https://camfi.readthedocs.io/. A PDF version of the documentation is provided at https://camfi.readthedocs.io/_/downloads/en/latest/pdf/.

### 2.1 Image collection

A total of ten wildlife cameras (BlazeVideo, model SL112) were mounted in various orientations at an alpine boulder field near Cabramurra, NSW (35°57’03S 148°23’50E, circa 1520 m elevation). The location was chosen as it is a known stopover point for Bogong moths on their forward migration. During the study period, light trapping was also being done in the area as part of other work, and an overwhelming majority of insects caught in these traps each night were Bogong moths (Linda Broome, pers. comm.). This fact assists our analysis by validating the assumption that most moths observed by the cameras were a single species (i.e. Bogong moths). The locations of four of the cameras are shown in Fig. 1a.

**Figure 1:**
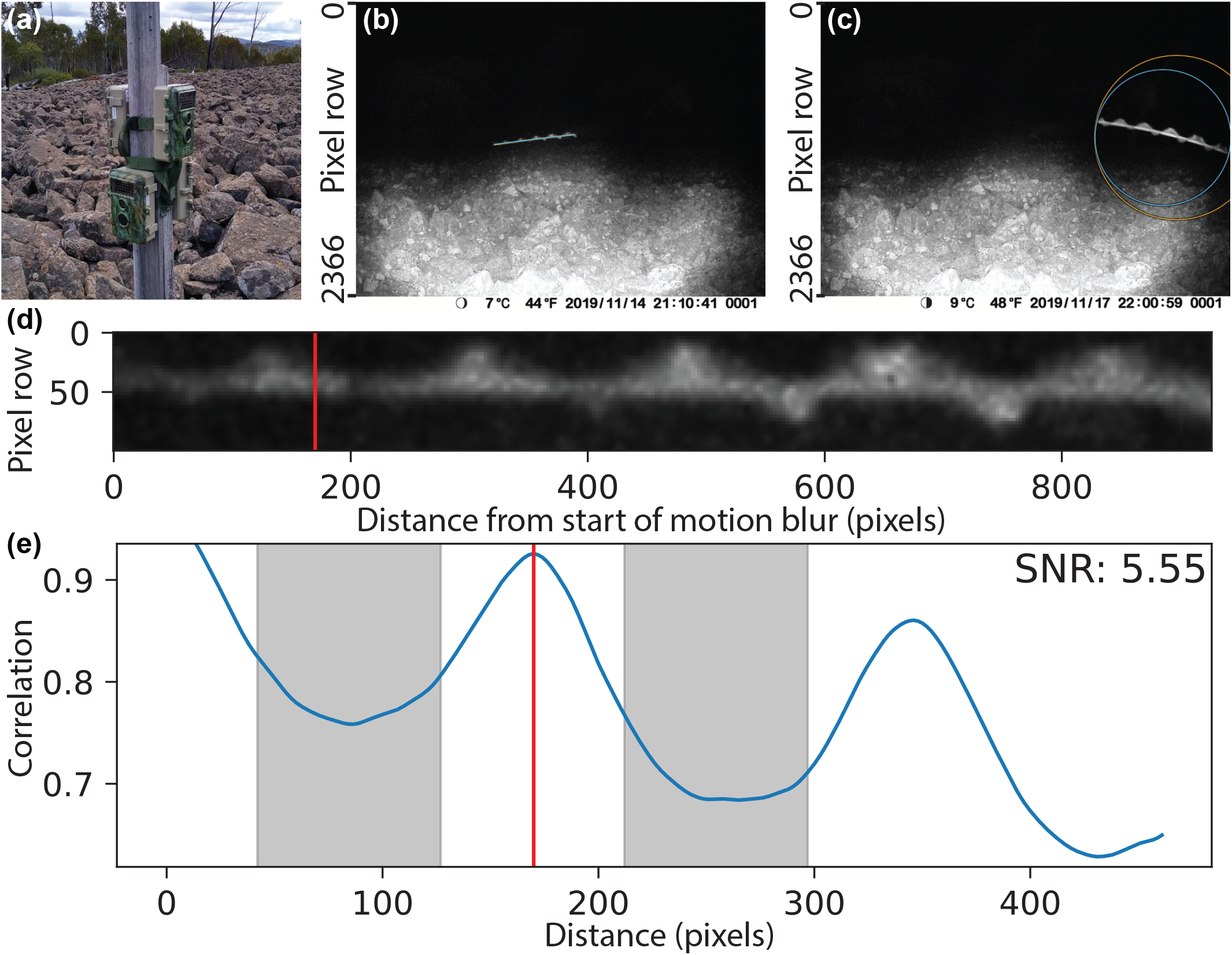
Example images showing data collection procedures used in this study. **(a)** Ten wildlife cameras (BlazeVideo, model SL112) were set to capture still photos on timers, and were deployed at the study site in a boulder field near Cabramurra, NSW in November 2019 (four cameras shown). **(b)** Motion blurs of moths captured by the cameras were marked with a polyline annotation. Manual annotation made in VIA (Dutta & Zisserman, 2019) is shown in *orange*, and the annotation made by our automated procedure is shown in *blue* (although since both annotations are very similar, they overlap and only the *blue* annotation is visible). **(c)** Circular or point annotations were used for images of moths whose motion blurs were not fully contained within the frame of the camera, or where the length of the motion blur was too short to see the moth’s wingbeat (latter case not shown). Manual annotation made in VIA (Dutta & Zisserman, 2019) is shown in *orange*, and the annotation made by our automated procedure is shown in *blue*. **(d)** Straightened and cropped “region-of-interest image”of moth motion blur, taken from image shown in b. Red vertical line shows periodicity along the axis of the motion blur as calculated by our algorithm. **(e)** Autocorellation of region-of-interest image (shown in d) along the axis of the motion blur. *Red line* shows peak periodicity as calculated by our algorithm. Signal-to-noise ratio (SNR) is calculated as the Z-score of the correlation at the peak, if drawn from a normal distribution with mean and variance equal to those of the correlation values within the *shaded regions*, defined by the intervals 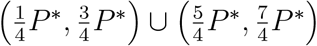, where *P*\* is the pixel period.

The cameras were set to capture photographs on a timer, capturing one photo every ten minutes, between the hours of 19:00 and 07:00 each night (Australian Eastern Daylight Time). Each camera was equipped with a 38-LED infra-red (940 nm wavelength) flash for unobtrusive night-time photography. Capture settings, such as ISO, exposure time, and whether to use the flash, were automatically selected by the camera before each capture, based on the ambient light levels. The cameras employ a fixed-focus lens.

The cameras were deployed on 14 November 2019 and collected on 26 November 2019, for a total of 11 nights of captures, resulting in a total of 8640 recorded images.

### 2.2 Image annotation

All images were manually annotated for flying moths using VIA (Dutta & Zisserman, 2019). It was noted that for many of the night-time shots, the exposure time was relatively long, which resulted in considerable motion blur from flying moths. In cases when this motion blur was completely contained within the frame of the camera, a polyline annotation from tip to tail of the motion blur was made in VIA, following the curved or straight path of the motion blur (illustrated in Fig. 1b). In cases where the motion blur was not completely within the frame of the camera, or where the motion blur was short with respect to the wingbeat, either a circular or point annotation was used instead (illustrated in Fig. 1c). Image metadata, including date and time of capture, and exposure time, were extracted from each of the images and incorporated into the output data file from VIA to enable downstream analyses, using our newly developed Python program, named “camfi.”

### 2.3 Automated annotation using Mask R-CNN

Although the process of manually annotating the images is simple to undertake, it is also time-consuming, particularly for large volumes of images. For large-scale studies, it may be desirable to use automated annotation, either by itself or in conjunction with manual annotation. To that end, we have developed an automatic annotation tool, which is included with camfi, and used by running `camfi annotate` from the command-line. The automatic annotation relies on Mask R-CNN (He et al., 2017), a state-of-the-art deep learning framework for object instance segmentation. The tool operates on VIA project files, allowing it to serve as a drop-in replacement for manual annotation. The tool also allows the annotations it generates to be loaded into VIA and manually edited if required.

#### 2.3.1 Training

To simplify training of the model to target other species, we have implemented a tool which automates the training process, and this is described below. This tool is packaged with camfi, and is used by running `camfi train` from the command-line. We have also included the model which we trained with camfi, so for species whose appearance is similar to that of Bogong moths while in flight, re-training the model may not be necessary.

We adopted the Mask R-CNN model architecture (He et al., 2017) with a Feature Pyramid Network (FPN) backbone (Lin et al., 2017). In the Mask R-CNN framework, the “head” of the model, which is the part of the model which generates the output, can include a variety of branches (corresponding to each type of output). We included the “class,” “bounding box,” and “instance segmentation” branches provided by the Mask R-CNN framework. We assigned two target classes; one for moths and another for background. We initialised the FPN backbone with a model which was pre-trained on the Common Objects in Context (COCO) object segmentation dataset (Lin et al., 2014) and employed the method of transfer learning to train the head and fine-tune the model. The data used for training were the set of images of flying moths we had previously annotated manually which contained at least one moth, as well as the corresponding manual annotations. This set contained 928 images (the remaining 7712 images from the full set of 8640 images had no moths in them). We reserved 50 randomly selected images from this set for testing, which were not seen by the model during training. Therefore, 878 images were used for training. The training data were augmented by creating new images by horizontal reflection of random individual images within the set of 878. In each iteration of training, a batch of five images^1^ were used. The model was trained for 15 epochs^1^ (full traversals of training data), for a total of 2640 iterations.

The manual annotations we used were polylines, points, and circles. However, Mask R-CNN operates on bounding boxes and segmentation masks, so some pre-processing of the annotations is required. These pre-processing steps are performed by our software directly on the output of the manual annotation process in VIA. For training the model, bounding boxes and segmentation masks are calculated on-the-fly from the coordinates of the manually annotated polylines, circles, and points. The bounding boxes are simply taken as the smallest bounding box of all coordinates in an annotation, plus a constant margin of ten pixels.^1^ The masks are produced by initialising a mask array with zeros, then setting the coordinates of the annotation in the mask array to one, followed by a morphological dilation of five pixels.^1^ For polyline annotations, all points along each of the line segments are set to one, whereas for point or circle annotations, just the pixel at the centre of the annotation is set.

We have made our annotation model available as part of the camfi software, and is the default model used by `camfi annotate`. We expect it to work out-of-the box for target species which are similar to the Bogong moth.

#### 2.3.2 Inference

Automation of the inference steps described in this section is implemented in the `camfi annotate` command-line tool, included with camfi. In inference mode, the Mask R-CNN model outputs candidate annotations for a given input image as a set of bounding boxes, class labels, segmentation masks (with a score from 0 to 1 for each pixel belonging to a particular object instance), and prediction scores (also from 0 to 1). Non-maximum suppression on candidate annotations is performed by calculating the weighted intersection over minimum (IoM) of segmentation masks of each pair of annotations in an image (the definition of IoM is provided in appendix A.1). For annotation pairs which have an IoM above 0.4,^2^ the annotation with the lower prediction score is removed. This has the effect of removing annotations which are too similar to each other, and are likely to relate to the same target. We also rejected candidate annotations with prediction scores below a given threshold.^2^ For each of the remaining candidate annotations, we fit a polyline annotation using the method described below.

To fit a polyline to a candidate annotation predicted by the Mask R-CNN model, we first perform a second-order^2^ polynomial regression on the coordinates of each pixel within the bounding box, with weights taken from the segmentation mask. If the bounding box is taller than it is wide, we take the row (y) coordinates of the pixels to be the independent variable for the regression, rather than the default column (x) coordinates. We then set the endpoints of the motion blur as the two points on the regression curve which lie within the bounding box, and which have an independent variable coordinate ten pixels^2^ away from the edges of the bounding box. The rationale for setting these points as the end points is that the model was trained to produce bounding boxes with a ten-pixel margin from the manual polyline annotations (see above). The curve is then approximated by a piecewise linear function (a polyline) by taking evenly spaced breakpoints along the curve such that change in angle between two adjoining line segments is no greater than approximately 15°.^2^

Finally, a check is performed on the polyline annotation to determine if the motion blur it represents is completely contained within the image. If it is not, it is converted to a circle annotation by calculating the smallest enclosing circle of all the points in the polyline annotation using Welzl’s algorithm (Welzl, 1991). The check is performed by measuring how close the annotation is to the edge of the image. If the annotation goes within 20 pixels^2^ of the edge of the image then the motion blur is considered to not be completely contained within the image, and therefore the polyline annotation is converted to a circle annotation.

The automatically produced annotations are saved to a VIA project file, and tagged with their prediction score, enabling further downstream filtering or annotation visualisation and diagnostics, as well as editing by a human if desired. We ran automatic annotation on the entire image set (8640 images) on a laptop with a Nvidia Quadro T2000 GPU. Using the GPU for inference is preferred, since it is much faster than using the CPU. However, in some cases, images which had a lot of moths in them could not be processed on the GPU due to memory constraints. To solve this problem, `camfi annotate` provides an option to run inference in a hybrid mode, which falls back to the CPU for images which fail on the GPU.

#### 2.3.3 Validation

This section introduces a number of terms which may be unfamiliar to the reader. Definitions of the following terms are provided in appendix A: intersection over union (appendix A.2), Hausdorff distance (appendix A.3), signed length difference (appendix A.4), precision-recall curve (appendix A.5), and average precision (appendix A.6). As mentioned above, we kept 50 randomly-selected annotated images as a test set during model training. We ran inference and validation on the full set of images, and on the test set in isolation. For both sets, we matched automatic annotations to the ground-truth manual annotations using a bounding-box intersection over union (IoU) threshold of 0.5.^3^ For each pair (automatic and ground-truth) of matched annotations we calculated IoU, and if both annotations were polyline annotations, we also calculated the Hausdorff distance *d_H_* and the signed length difference Δ*L* between the two annotations. Gaussian kernel density estimates of prediction score versus each of these metrics were plotted for diagnostic purposes. We also plotted the precision-recall curve and calculated the average precision *AP*_50_ for both image sets. To compare future automatic annotation methods to ours, we recommend using mean IoU 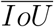, mean Hausdorff distance 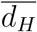, mean length difference 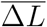, the standard deviation of length difference *σ*_Δ*L*_, and *AF*_50_ as the set of comparison metrics.

### 2.4 Wingbeat frequency measurement

For observations of moths whose motion blur was entirely captured within the frame of a camera, we use a polyline annotation, which follows the path of the motion blur. This annotation can be obtained either manually or automatically, by the procedures described above. For the analyses presented in this paper, we used manual annotations. Since the moth is moving while beating its wings, we are able to observe the moth’s wingbeat (see Fig. 1b). Incorporating information about the exposure time and rolling shutter rate of the camera, we are able to make a measurement of the moth’s wingbeat frequency in hertz. We have implemented the procedure for making this measurement as part of camfi, in the sub-command called `camfi extract-wingbeats`. The procedure takes images from wildlife cameras (like those shown in Fig. 1b,c) and a VIA project file containing polyline annotations of flying insect motion blurs as input, and outputs estimates of wingbeat frequencies and other related measurements. A description of the procedure for a given motion blur annotation follows.

First, a region of interest image of the motion blur is extracted from the photograph, which contains a straightened copy of the motion blur only (see Fig. 1d). The precise method for generating this region of interest image is not important, provided it does not scale the motion blur, particularly in the direction of the motion blur. Our implementation simply concatenates the rotated and cropped image rectangles, which are centred on each segment of the polylines, with length equal to the respective segment, and with an arbitrary fixed width.^4^ We used the default value of 100 pixels.

The pixel-period of the wingbeat, which we denote *P*, is determined from the region of interest image by finding peaks in the autocorrelation of the image along the axis of the motion blur (see Fig. 1e). The signal-to-noise ratio (SNR) of each peak is estimated by taking the Z-score of the correlation at the peak, if drawn from a normal distribution with mean and variance equal to those of the correlation values within the regions defined by the intervals 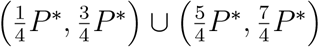, where *P*\* is the pixel period corresponding to the given peak. The peak with the highest SNR is selected as corresponding to the wingbeat of the moth and is assigned as *P*. The total length of the motion blur (in pixels) may then be divided by *P* to obtain a non-integer wingbeat count for the motion blur. The SNR of the best peak is included in the output of the program, to allow for filtering of wingbeat data points with low SNR. It should be noted that the definition of SNR used here may differ somewhat from other formal definitions. For example, this definition admits negative values for SNR (albeit rarely), in which case the corresponding measurement will surely be filtered out after a SNR threshold is applied. When running `camfi extract-wingbeats`, supplementary figures containing the region of interest images and corresponding autocorrelation plots, similar to those presented in Fig. 1d,e can be optionally generated for every polyline annotation.

To calculate wingbeat frequency *F_w_*, in hertz, we need to know the length of time that the moth was exposed to the camera, which we call Δ*t*. Unfortunately, this is not as simple as taking the exposure time as reported by the camera, which we call *t_e_*, due to the interaction of the first-order motion of the moth with the rolling shutter of the camera. In particular,

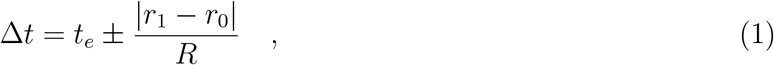

where *r*_0_ and *r*_1_ are the row indices (counting rows from the top of the image) of the two respective ends of the polyline annotation, and *R* is the rolling shutter line rate, which was measured to be 9.05 × 10^4^ lines *s*^−1^ for the cameras we used (for a method of measuring *R*, see appendix A.7). The “±” reflects the fact that it is impossible to tell in which direction the moth is flying from the images alone, leading us to two possible measurements of moth exposure time, corresponding to the moth flying down or up within the image plane of the camera, respectively. Under certain circumstances, this ambiguity can be resolved by observing that Δ*t* ≥ 0, i.e. insects cannot fly backwards through time. Intuitively, we may then attempt to calculate *F_w_* by dividing the wingbeat count by Δ*t* (these preliminary estimates of wingbeat frequency are included in the output of `camfi extract-wingbeats`). However, this would require the assumption that the moth has a body length of zero, since the length of the motion blur, which we denote as *L*, is the sum of the first order motion of the moth during the exposure and the moth’s body length, projected onto the plane of the camera. Clearly, this assumption may be violated, as the insects have a non-zero body length in the images. We denote the body length of the moth projected onto the plane of the camera by the random variable *L_b_*.

The statistical procedure for estimating the mean and standard deviation of observed moth wingbeat frequency, which accounts for both the time ambiguity and the non-zero body lengths of the moths, is as follows. We begin with the following model, which relates *F_w_* to *L_b_* and various measured variables.

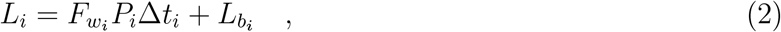

where *i* is the index of the observation. We proceed by performing a linear regression of *L* on *P*Δ*t* (setting *P*Δ*t* as the independent variable) using the BCES method (Akritas & Bershady, 1996) to obtain unbiased estimators of 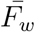 and 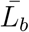, as well as their respective variances, 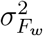 and 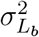. Values for Δ*t_i_* are taken as the midpoints of the pairs calculated in equation 1, with error terms equal to 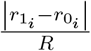. Values for *L_i_* are assumed to have no measurement error. Where multiple species with different characteristic wingbeat frequencies are observed, an expectation-maximisation (EM) algorithm may be applied to classify measurements into groups which may then be analysed separately. We may then test the zero body length assumption, namely 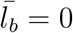, by calculating its *t* statistic.

### 2.5 Implementation

Our implementation of camfi and its associated tools is written in Python 3.9 (Python Software Foundation, https://www.python.org/). The latest version of camfi relies on (in alphabetical order): bces 1.0.3 (Nemmen et al., 2012), exif 1.3.1 (Thieding et al., 2021), imageio 2.9.0 (Silvester et al., 2020), Jupyter 1.0.0 (Kluyver et al., 2016), Matplotlib 3.4.2 (Hunter, 2007), NumPy 1.21.1 (Harris et al., 2020), Pandas 1.3.0 (McKinney et al., 2010), Pillow 8.3.1 (Kemenade et al., 2021), pydantic 1.8.2 (Colvin et al., 2021), Scikit-image 0.18.2 (Van der Walt et al., 2014), Scikit-learn 0.24.2 (Pedregosa et al., 2011), SciPy 1.7.0 (Virtanen et al., 2020), Shapely 1.7.1 (Gillies et al., 2007--), skyfield 1.39 (Rhodes, 2019), Statsmodels 0.12.2 (Seabold & Perktold, 2010), strictyaml 1.4.4, PyTorch 1.9.0 (Paszke et al., 2019), TorchVision 0.10.0 (Marcel & Rodriguez, 2010), and tqdm 4.61.2 (Costa-Luis et al., 2021).

Camfi is open source and available under the MIT license. The full source code for the latest version of camfi and all analyses presented in this paper are provided at https://github.com/J-Wall/camfi. The documentation for camfi is provided at https://camfi.readthedocs.io/. Camfi is under active development and we expect new features and new trained models to be added as new versions of camfi are released from time to time. All analyses presented in this paper were done using camfi 2.1.3, which is permanently available from the Zenodo repository https://doi.org/10.5281/zenodo.5194496 (Wallace, 2021b).

## 3 Results

### 3.1 Moth activity patterns

From the 8640 images analysed, a total of 1419 manual annotations were made. Of these, 259 were circle or point annotations, which we are able to use for quantifying general activity, but which cannot be used for wingbeat analysis. The remaining 1160 annotations were polyline annotations, which we used for both activity quantification and wingbeat analysis.

We observed a daily pattern of moth activity, with marked increase in the number of moths flying during evening twilight on most days (Fig. 2a). This daily pattern is clearly pronounced in Fig. 2b. By considering just the period of evening twilight from each day of the study, we are able to quantify the relative evening moth activity levels over time, and compare them with abiotic factors such as the weather (Fig. 2c). We performed a Poisson regression of evening twilight moth activity levels (number of annotations, with exposure set to the number of images taken during evening twilight that day) against daily weather factors, (Bureau of Meteorology, 2019) and found that minimum morning temperature, minimum evening temperature, daylight hours, and temperature range had a significant joint effect on observed moth numbers (Fig. 2c, green trace: Wald test, *χ*^2^ = 25.3, *df* = 4, *p* ≪ 0.001).

**Figure 2:**
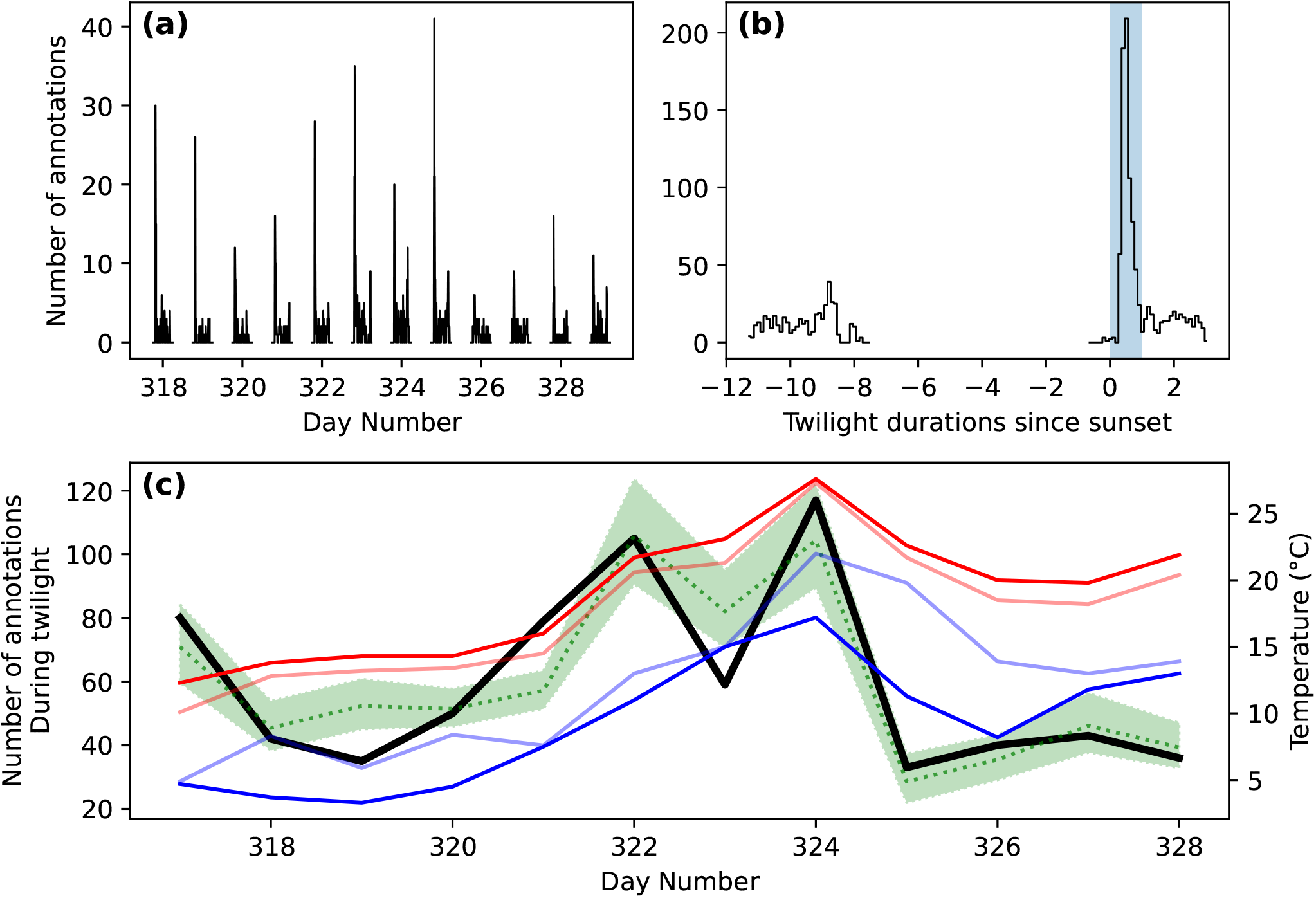
Moth activity levels during the November study period. **(a)** Number of moths observed across the study period from 10 cameras, with images taken at 10 min intervals. **(b)** Total number of moth observations by time after sunset (scaled by the duration of twilight) shows peak in activity during evening twilight (shaded blue). **(c)** Number of moths observed during twilight for each day of the study (black), shown with daily temperatures recorded by the BOM at nearby Cabramurra (Bureau of Meteorology, 2019): maximum (red), minimum (blue), 9 am (light blue), and 3 pm (light red). Predicted values for moth activity (and SE confidence interval) from a Poisson regression of number of annotations vs. daily weather factors (minimum morning temperature, minimum evening temperature, daylight hours, and temperature range) are shown in green.

### 3.2 Wingbeat frequency

Of the 1160 manual polyline annotations of flying moths, 580 yielded wingbeat measurements which had a SNR exceeding the threshold of 4.0 (Fig. 3a). The histogram of preliminary wingbeat calculations, which do not account for non-zero body-length (Fig. 3a), indicated that there were likely to be two classes of insect wingbeats observed, possibly corresponding to two separate species. It was noted that the preliminary wingbeat frequencies of the less common of the two classes were centred at a value approximately half that of the more common class, indicating the possibility that the lesser class represented an artefact of the wingbeat measurement process, where a given signal’s period could conceivably be inadvertently doubled. To rule out this possibility, a subset of the wingbeat region-of-interest images were viewed, and no obvious evidence of erroneous measurements was observed. It was therefore concluded that the two observed classes of wingbeats represented a true biological signal, so for subsequent analysis we assumed there are two types of wingbeat represented in the data.

**Figure 3:**
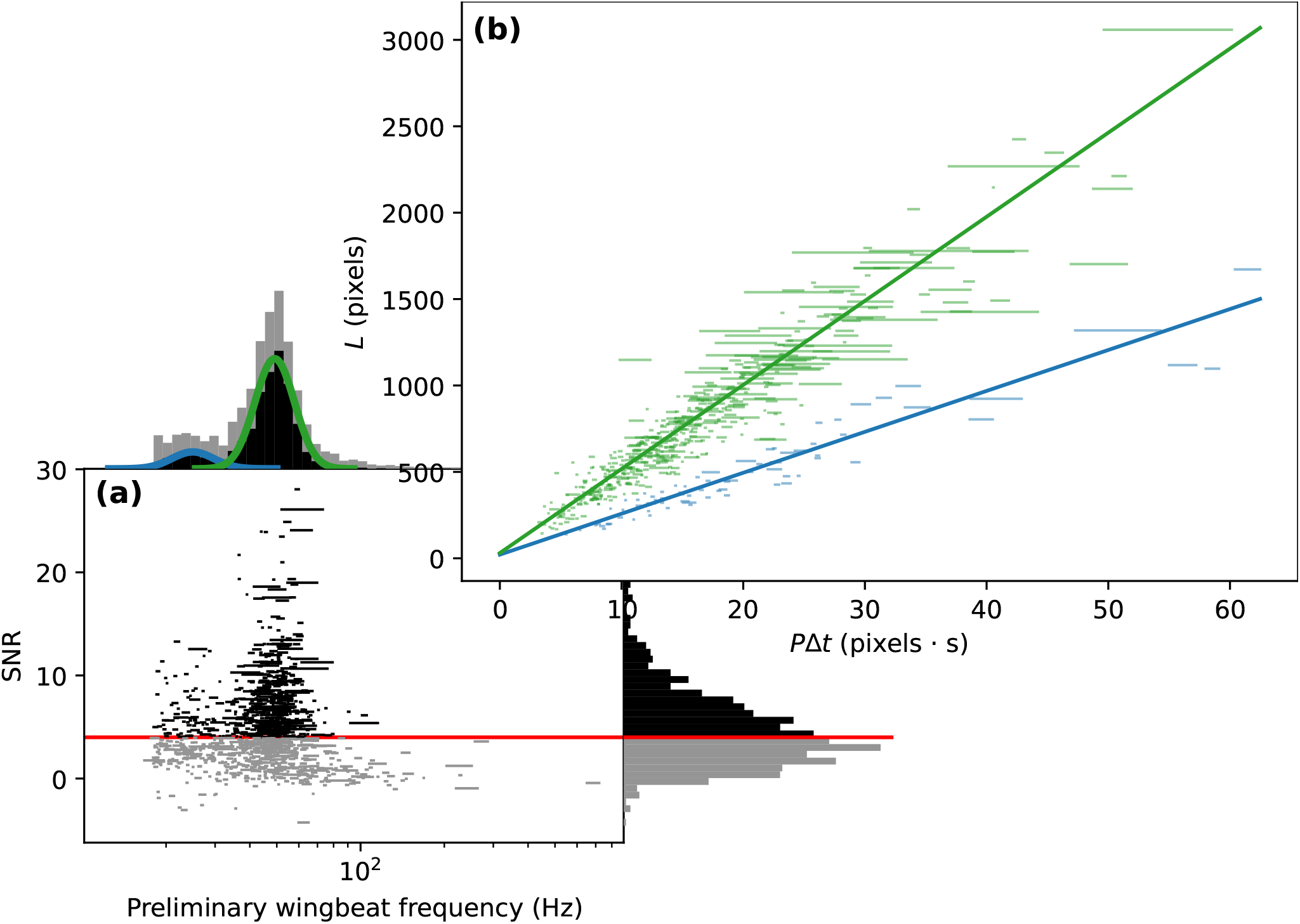
Moth wingbeat frequency measurements from wildlife camera images. Error bars indicate the two possible measurements arising from each observation, due to the interaction between flightdirection ambiguity and the rolling shutter of the cameras (equation 1). **(a)** Signal-to-noise ratio (SNR) vs. preliminary wingbeat frequency measurements on log_10_ scale, with SNR threshold (4.0) indicated by red line. Preliminary wingbeat measurements do not account for non-zero body length of observed moths. Marginal distribution histograms for both axes are shown. Data which exceeded the SNR threshold are in dark grey, and data which did not meet the SNR threshold are in light grey. The probability density functions associated with a log_10_-gaussian mixture model (GMM) of above-threshold preliminary wingbeat frequencies for two target classes are overlaid on the horizontal marginal distribution histogram (blue and green curves). **(b)** Length of motion-blur *L* vs. pixelwingbeat period × exposure time of motion-blur *P*Δ*t* for observations which exceeded SNR threshold. Linear regressions are shown, which were obtained by the BCES method (Akritas & Bershady, 1996), extended to classify the data into two target classes, and regress each class separately using an expectation-maximisation (EM) algorithm. This regression eliminates the assumption of zero-body length. For convenience, the two classes are coloured in the same way as in panel a, however it should be noted that the classifications presented in each sub-figure are distinct. The slopes of the regressions estimate mean wingbeat frequency, and the intercepts estimate mean non-zero pixel-body length.

To produce unbiased estimates of mean wingbeat frequency (which accounts for non-zero body-length), we performed a linear regression of *L* vs. *P*Δ*t* using the BCES method (Akritas & Bershady, 1996) (Fig. 3b). Two target classes were identified using an EM algorithm, and regressed separately. The EM algorithm assigned 75 observations to the first class, and 505 observations to the second class, which we infer as representing Bogong moths. The slopes of the linear regressions give estimates of mean wingbeat frequency, and these are 23.7 Hz (SE = 1.8) and 48.6 Hz (SE = 1.4) for the two classes, respectively. The intercepts give estimates of mean pixel-body lengths at 21.2 pixels (SE = 30.4) and 30.4 pixels (SE = 18.9). The *t* statistics of the body length estimates are 0.6972 and 1.6067 respectively, for the null hypothesis that body length is zero. This leads to one-sided p-values of 0.244 and 0.054, respectively. Since both of these are above the canonical p-value threshold of 0.05, we conclude that the zero body length assumption is reasonable, and that a log-gaussian mixture model is sufficient to describe the observed wingbeat frequencies. After correcting for the (known) measurement error produced by the interaction between flight-direction ambiguity and the rolling shutter of the cameras (equation 1), The log-gaussian mixture model gives us estimates of wingbeat frequencies, which are 25.10 Hz (std = 2.88) and 49.40 Hz (std = 5.25), for the two classes of moth observed, respectively.

### 3.3 Automatic annotation

Automatic annotation performance was evaluated using a test set of 50 images, as well as the full set of 8640 images. Evaluation metrics for both sets are presented in table 1. Each metric was similar across both image sets, indicating that the annotation model has not suffered from over-fitting. This is also supported by the contour plots of prediction score vs. IoU, polyline Hausdorff distance, and polyline length difference (Fig. 4b,c,d, respectively). These plots show similar performance on both the full image set (8640 images) and the test set (50 images). Furthermore, they show that prediction scores for matched annotations (automatic annotations which were successfully matched to annotations in the manual ground-truth dataset) tended to be quite high, as did the IoU of those annotations, while both polyline Hausdorff distance *d_H_* and polyline length difference Δ*L* clustered relatively close to zero. The precision-recall curves of the automatic annotator (Fig. 4e) show similar performance between the image sets, and show a drop in precision for recall values above 0.6. Training took 2640 iterations and completed in less than 2 h (Fig. 4a) on a machine with two 8-core Intel Xeon E5-2660 CPUs running at 2.2GHz and a Nvidia T4 GPU, and inference took on average 1.15 s per image on a laptop with a 6-core Intel Xeon E-2276M CPU running at 2.8GHz and a Nvidia Quadro T2000 GPU.

**Table 1:** Automatic annotation performance metrics when tested against the full image set (8640 images), and the test set (50 images). Performance metrics calculated are average precision *AP*_50_, mean bounding-box intersection over union 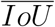, mean Hausdorff distance of polyline annotations 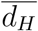, mean signed length difference of polyline annotations 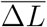, and the standard deviation of signed length difference of polyline annotations *σ*_Δ*L*_. Definitions of these metrics are provided in appendix A.

| Image set | $AP_{50}$ | $\overline{IoU}$ | $\overline{d_H}$ | $\overline{\Delta L}$ | $\sigma_{\Delta L}$ |
| --- | --- | --- | --- | --- | --- |
| Full set | 0.588 | 0.814 | 29.2 | -3.31 | 46.4 |
| Test set | 0.687 | 0.805 | 28.8 | 5.16 | 51.0 |

**Figure 4:**
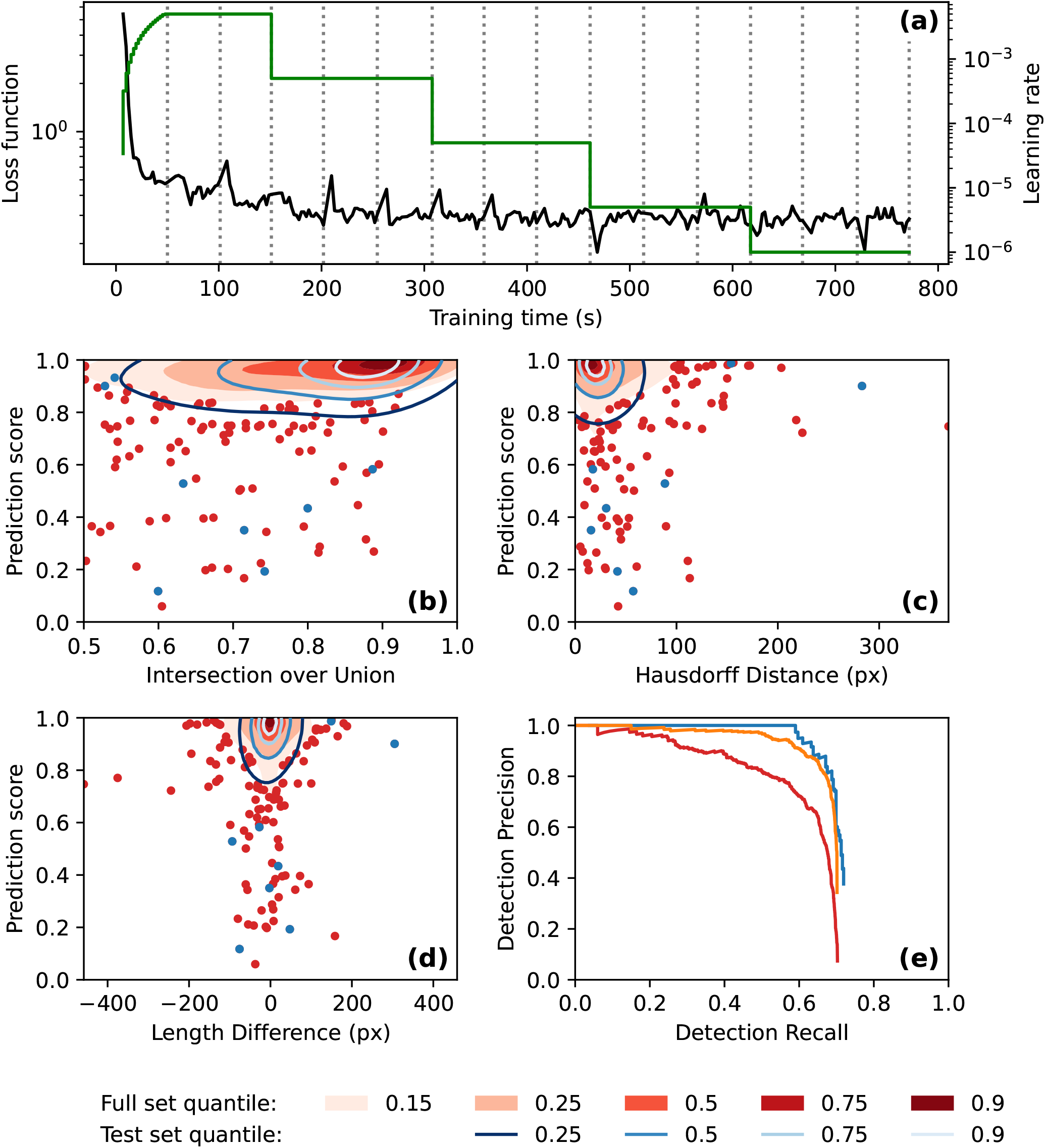
Automatic annotation evaluation plots. **(a)** Automatic annotation model training learning rate schedule (green) and loss function (black) over the course of training. Epochs (complete training data traversal) are shown with dotted vertical lines. **(b)-(e)** Similar performance was seen for both the full 8640-image set (red) and the test 50-image set (blue). **(b)-(d)** Gaussian kernel density estimate contour plots of prediction score vs. **(b)** bounding box intersection over union, **(c)** polyline Hausdorff distance, and **(d)** polyline length difference, for both image sets. Contours are coloured according to density quantile (key at bottom of figure). In each plot, data which lie outside of the lowest density quantile contour are displayed as points. **(e)** Motion blur detection precision-recall curve, generated by varying prediction score threshold. The precision-recall curve for the set of 928 images which had at least one manual annotation is shown in orange.

## 4 Discussion

This paper demonstrates the utility of inexpensive wildlife cameras for the long-term monitoring of activity in flying insects, and describes how they may be used to measure the wingbeat frequency of those insects. We do not expect this method to completely replace other approaches for monitoring insects, such as trapping, which enables precise measurement of biodiversity and positive identification of species. Likewise, it will not completely replace other remote sensing approaches, such as radar and lidar, which facilitate detecting targets at long distances. However, it is clear that this method has significant potential to complement these other approaches, and in certain circumstances, replace them. For instance, in comparison to these other approaches, this method is particularly suited to monitoring assemblages of known species in remote areas, especially when it is known that the target insects are low-flying. An advantage of the presented method over trapping is that much greater temporal resolution is gained. In the present study one measurement was taken by each camera every ten minutes, and depending on the research question or absolute abundance of the insects being studied, this can easily be varied. This is in contrast to trapping studies, where only one measurement of abundance can be recorded per visit to the trap by the researcher. This provides an opportunity to use the present method to answer a variety of ethological research questions which may not be approachable with previous methods.

The measurement of wingbeat frequency has utility in distinguishing between multiple target species, especially when the observed flying insects are dominated by one known species, as is the case for the dataset we analysed, or where the wingbeat frequencies of the observed species are very different from each other. The method of generating summary statistics for observed wingbeat frequency is complicated somewhat by the measurement error introduced by the interaction between the ambiguity in insect flight direction and the rolling shutter of the cameras. This measurement error could be eliminated in one of two ways: 1. By taking two immediately successive exposures, which would enable inference of flight direction, or 2. By using cameras with a global shutter, which would prevent the flight direction of the insect from having any influence over the duration that the insect is exposed to the camera. Implementing either of these options is desirable, however they are not possible without significantly more expensive cameras than the type used in this study. This would limit the utility of the method for use in either large-scale or low-budget studies. Until the cost of wildlife cameras equipped with a global-shutter comes down, the most practical approach remains to handle this measurement error statistically.

This paper has presented a method for monitoring nocturnal flying insects, however there is no reason it couldn’t be used for diurnal species as well, provided care is taken with regard to the placement of cameras. Namely, it would be important to have a relatively uniform background (such as the sky) in order to be able to see insects in the images during the day. In this case, the infra-red flash of the cameras would not be used and the insects would appear as dark objects on a light background. During the day, the exposure time of the cameras is much shorter than at night, so it would be impossible to use this method to measure wingbeat frequencies of day-flying insects. However, in some cases it may be possible to identify day-flying insects in the images directly. It may also be possible to recreate the type of images seen during the night in any lighting conditions by retrofitting the cameras with long-pass infra-red filters, neutral density filters, or a combination of both.

A key advantage of the present method over other approaches is that it can be readily scaled to large monitoring studies or programs, thanks to the low cost of implementation and the inclusion of the tool for automatic annotation of flying insect motion blurs. It is expected that studies implementing this method for target species which substantially differ in appearance from Bogong moths when in flight (and where the use of automatic annotation is desired) may have to re-train the Mask R-CNN instance segmentation model. We believe that the tools we have implemented make that process highly accessible.

## 5 Acknowledgements

EJW and JRAW are grateful for funding from the European Research Council (Advanced Grant No. 41298 to EJW), and the Royal Physiographic Society of Lund (to JRAW). JRAW is thankful for the support of an Australian Government Research Training Program (RTP) Scholarship. We thank Drs. Jochen Zeil, Ryszard Maleszka, Linda Broome, Ken Green, Samuel Jansson, Alistair Drake, Benjamin Amenuvegbe, Mr. Benjamin Mathews-Hunter, and Ms. Dzifa Amenuvegbe Wallace for invaluable collaboration and assistance, and Drs. John Clarke and Francis Hui for useful discussions relating to the wingbeat frequency analyses.

## 6 Author’s contributions

JRAW, DD and EJW conceived the ideas; JRAW, TR, BB and DD designed the methodology; JRAW and TR collected the data; JRAW and BB analysed the data; JRAW led the writing of the manuscript with significant input from EJW. All authors contributed critically to the drafts and gave final approval for publication.

## 7 Data Availability

The images and manual annotations are available from the Zenodo repository https://doi.org/10.5281/zenodo.4950570 (Wallace, 2021a). All other data and code are available from the Zenodo repository https://doi.org/10.5281/zenodo.5194496 (Wallace, 2021b).

## A Appendix

### A.1 Weighted Intersection over Minimum (IoM)

It is common for automatic image annotation procedures to produce multiple candidate annotations for a single object of interest (in our case, motion blurs of flying insects). It is therefore necessary to perform non-maximum suppression on the automatically generated candidate annotations (where only the best candidate annotation for each object is kept).

In order to perform non-maximum suppression on candidate annotations, we need a way of matching annotations which refer to the same object. This is typically done by defining some measure of similarity between two annotations, and then applying this measure to each pair of annotations within an image. Then, by setting an appropriate threshold, the program can decide which pairs of annotations require non-maximum suppression to be applied.

In the case of our method, it is common for the automatic annotation procedure to produce multiple annotations of different sizes for each motion blur. This is likely due to the fact that the motion blurs themselves can vary greatly in length and in number of wingbeats, which causes some level of confusion for the automatic annotation model. Therefore, we need to use a similarity measure which is invariant to the size of annotations, and produces a high similarity for annotations which are (roughly) contained within each other. This motivates our definition of similarity of candidate annotations for the purposes of non-maximum suppression. Namely, weighted intersection over minimum (IoM).

Suppose we have two sets *A* = {*a*_0_, *a*_1_, …, *a_n_*} and *B* = {*b*_0_, *b*_1_,…, *b_m_*} with corresponding weights *X* = {*x*(*a*_0_), *x*(*a*_1_),…, *x*(*a_n_*)} and *Y* = {*y*(*b*_0_), *y*(*b*_1_), …, *y*(*b_m_*)}. We define the intersection over minimum of the two sets as

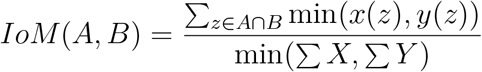

In camfi, we apply this definition by setting the weights *X* and *Y* as the segmentation mask values from two candidate annotations respectively. In this case, *A* = *B* are the coordinates of every pixel in the image.

### A.2 Bounding-box Intersection over Union (IoU)

To validate the quality of an automatic annotation system, we would like to compare the annotations produced by the system to annotations produced by a human. To do this, we need to have a way of matching pairs of annotations. This can be done by measuring the similarity between two annotations, and if they are similar enough, matching them.

The bounding-box intersection over union (IoU) is a commonly used similarity measure for object detection on images. It is defined as per it’s name. That is, we find the bounding box of two annotations, then calculate the ratio of the intersection of the two boxes with the union of the two boxes.

The mean bounding-box intersection over union 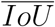 is the arithmetic mean of all *IoU* values across the automatic annotations which were successfully matched to a manual ground-truth annotation. Since matches were made for annotations with *IoU* > 0.5, it must also hold that 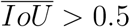.

### A.3 Hausdorff distance

Our method for measuring wingbeat frequencies depends on accurate annotations of flying insect motion blurs, so it is important to know the accuracy of the annotations produced by our method for automatic annotation.

Suppose we have an automatically generated polyline annotation, and a corresponding polyline annotation made by a human which we would like to validate the automatic annotation against. We would like to know how accurately the automatic annotation recreates the human annotation. We proceed by calculating the Hausdorff distance between the two annotations. First, we define two sets *A* and *B* which contain all the points on the respective polyline from each of the two annotations. The Hausdorff distance *d_H_*(*A, B*) is defined as

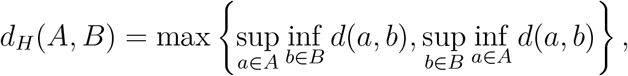

where sup is the supremum, inf is the infimum, and *d*(*a, b*) is the Euclidean distance between points *a* and *b*. In other words, the Hausdorff distance is the maximum distance between a point in one of the sets, to the closest point in the other. For the purpose of validating automatic annotations, we see that smaller Hausdorff distances between the automatic and manual annotations are better than larger ones.

The mean Hausdorff distance 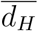 is the arithmetic mean of all values of *d_H_* across the automatic polyline annotations which were successfully matched to polyline annotations in the manual ground-truth dataset.

### A.4 Signed Length Difference

Another way to assess the accuracy of the automatic polyline annotations against the manually produced annotations is signed length difference Δ*L*. This is motivated by the fact that our method for calculating wingbeat frequency is fairly sensitive to the length of the polyline annotation. Suppose we have an automatically generated polyline annotation with length *L_A_* and a corresponding ground-truth manual annotation with length *L_G_*. Then the signed length difference is defined as Δ*L* = *L_A_* − *L_G_*. The closer the signed length difference is to zero, the better.

The mean signed length difference 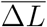 is the arithmetic mean of all values of Δ*L* across the automatic polyline annotations which were successfully matched to polyline annotations in the manual groundtruth dataset. The standard deviation of signed length difference *σ*_Δ*L*_ is the standard deviation of these values.

### A.5 Precision-Recall curve

With regard to object detection, precision is the proportion of detections which correspond to annotations present in the ground-truth dataset. Recall is the proportion of objects in the ground-truth dataset which are detected by the automatic annotation system. In our case, the ground-truth dataset is the set of manual annotations. We match automatic annotations with ground-truth annotations if they have an IoU greater than 0.5.

Each candidate annotation is given a confidence score between 0.0 and 1.0 by the annotation model. This score can be used to filter the candidate annotations (e.g. by removing all annotations with a score less than 0.9). By varying the score threshold, we obtain different precision and recall values for the system.

A precision-recall curve is the curve drawn on a plot of precision vs. recall by varying the score threshold. The closer the curve goes towards the point (1, 1), the better.

### A.6 Average precision

The average precision *AP*_50_ is calculated from the precision-recall curve. It is simply the average (arithmetic mean) of the precision values at the following recall values: 0.0, 0.1, 0.2, 0.3, 0.4, 0.5, 0.6, 0.7, 0.8, 0.9, and 1.0.

### A.7 Measurement of rolling shutter line rate

For the purposes of measuring the wingbeat frequency of the moths in the images captured by the wildlife cameras, it is important to know the line rate of the cameras’ rolling shutters. This was measured by mounting one of the cameras so that its lens pointed at a rotating white line (in this case, a strip of paper taped to a cardboard tube attached to the blades of a small electric fan), ensuring that the centre of the white line and its centre of rotation were coincident. The apparatus for measuring the rolling shutter line rate of the cameras is shown in Fig. 5.

**Figure 5:**
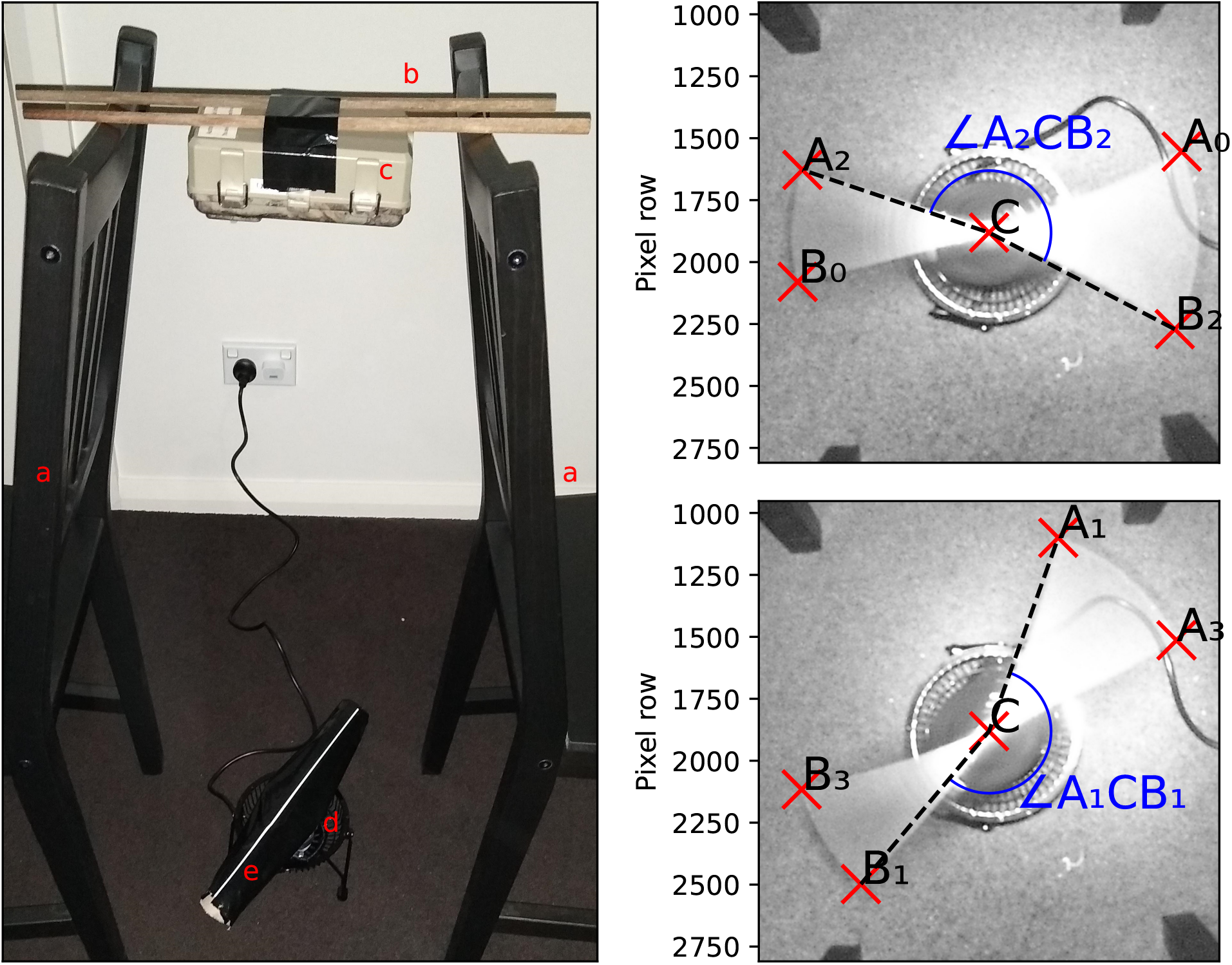
Measurement of rolling shutter line rate of the wildlife cameras used in this study. **Left:** Apparatus for measuring rolling shutter line rate. **a.** Camera supports (chairs). **b.** Camera mount (long chopsticks with duct tape). **c.** Camera to be measured. **d.** Rotor motor (desk fan with front cover removed). **e.** Rotor (cardboard tube) with white reference line (paper masked with black duct tape). **Right:** Two rolling shutter line rate measurement images, with reference annotations marked, and angles ∠*A*_1_*CB*_1_ and ∠*A*_2_*CB*_2_ shown. Using the coordinates of these reference annotations, and if the rotational velocity of the white reference line is known, the rolling shutter line rate of the camera can be calculated using equation 3.

The exact rotational velocity of the line was measured by synchronising a strobe light from a smart phone application (Strobily, 2019) to the period of rotation of the line. Synchronisation is achieved when the line appears stationary under the strobe.

A photograph was taken using the camera and the corners of the motion blur traced by the rotation line were marked using the free and open-source VGG Image Annotator (VIA) (Dutta & Zisserman, 2019) (VIA is a simple and standalone manual annotation software for image, audio and video, and is available from https://www.robots.ox.ac.uk/~vgg/software/via/). The marked corners correspond to the positions where the exposure of the rotating line began and ended (see Fig. 5). Since we know the rotational velocity of the line, we can then calculate the rolling shutter line rate from the coordinates of these corners. Namely,

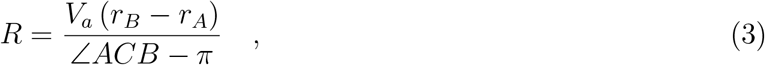

where *R* is the rolling shutter line rate, *V_a_* is the rotational velocity of the white line (in rad *s*^−1^), *A* and *B* are the coordinates of two opposite corners of the motion blur traced by the rotating line (with A being the corner with the lower pixel-row index), *C* is the centre of rotation of the line, and *r_A_* and *r_B_* are the pixel-row indices of *A* and *B* respectively. Following the convention for digital images, the first pixel-row is at the top of the image (see Fig. 5).

Rolling shutter line rate was calculated for both the start of exposure (Fig. 5: *A*_0_ and *B*_0_, *A*_1_ and *B*_1_) and end of exposure (Fig. 5: *A*_2_ and *B*_2_, *A*_3_ and *B*_3_), for two separate images. These four calculated values of *R* were very similar, so it was assumed that this particular model of camera has one constant rolling shutter line rate. This rate was taken to be the average of the two measured values from the second image, or 9.05 × 10^4^ lines *s*^−1^, which was used for all subsequent analyses, since only one model of camera was used in this study. If a different model is used, we would recommend repeating this measurement.

## Footnotes

1 Configurable parameter when running Camfi.

2 Configurable parameter when running Camfi.

3 Configurable parameter when running Camfi.

4 Configurable parameter when running Camfi.

